# Drought impact on phellem development: identification of novel gene regulators and evidence of photosynthetic activity

**DOI:** 10.1101/2023.12.26.573371

**Authors:** Pedro M. Barros, Helena Sapeta, Diogo A. Lucas, M. Margarida Oliveira

## Abstract

*Quercus suber* (cork oak) is a sustainably exploited forest resource, producing a unique renewable raw material known as cork. With drought events imposing a negative impact on tree vitality, we need more knowledge on the genetic and environmental regulation of cork development to protect the cork sector. We focused on characterizing long-term drought-induced molecular adaptations occurring in stems, and identifying key genetic pathways regulating phellem development. One-year-old cork oak plants were grown for 6 months under well-watered, or water-deficit (WD) conditions and main stems were targeted for histological characterization and transcriptomic analysis. WD treatment impaired secondary growth, by reducing meristem activity at both vascular cambium and phellogen. We analyzed the transcriptional changes imposed by WD in phellem, inner bark, and xylem, and found a global downregulation of genes related to cell division, differentiation, and cell wall biogenesis. Phellem and inner bark showed upregulation of photosynthesis-related genes, highlighting a determinant role of stem photosynthesis in the adaptation to long-term drought. We show that developing phellem cells contain chloroplasts and their abundance increases under WD. Finally, we propose new candidate regulatory genes involved in the regulation of phellogen activity and demonstrate the involvement of phellem in drought-induced bark photosynthesis in young plants.

**Highlight:** Phellem development in cork oak is impaired in drought adaptation, by negative regulation of cell division and differentiation programs, while photosynthesis is induced to contributing to CO_2_ recycling in the stem.

## Introduction

Cork is a natural and fully renewable forest product obtained from the bark of *Quercus suber* (cork oak), with high commercial interest given its unique physical and chemical properties (Pereira, 2007; Teixeira, 2022). Cork culture also plays an important ecological role by promoting the maintenance of the traditional farming landscapes (Pereira, 2007) and contributing to carbon sequestration in these regions (Costa-e-Silva *et al*., 2021).

While the commercial cork product is an almost exclusive feature of cork oak, the mechanisms leading to its formation are quite conserved in most plants displaying secondary growth, from herbaceous angiosperms to woody plants (reviewed by Serra *et al*., 2022). After the establishment of the vascular cambium and initiation of radial growth, there is the formation of a new meristem known as phellogen that starts dividing bifacially forming phelloderm cells inwards and phellem cells outwards. In stems, phellogen establishes in the sub-epidermal layer, replacing the epidermis at the plant-environment interface, and establishing the periderm. Phelloderm cells are parenchyma cells with similar morphology to underlying tissues from the inner bark (cortical parenchyma and phloem), likely playing similar functions, such as storage (Serra *et al*., 2022), and displaying photosynthetic activity (Gartner, 1996; Burrows and Connor, 2020). Phellem cells establish the outer bark by undergoing a process of cell differentiation that includes (i) the establishment of a specialized secondary cell wall enriched mostly in suberin, (ii) an increase in volume and (iii) the induction of programmed cell death. In cork oak, among other species, phellogen activity is imbalanced and generates mostly phellem (Graça and Pereira, 2004). The high activity and longevity of this cambium in cork oak leads to the formation of a homogeneous layer of phellem, that is harvested by the physical (manual) rupture of active phellogen cells.

In recent years, the molecular dynamics driving phellem development has received increased attention. In cork oak, different genome-wide transcriptomic studies have described distinctive transcriptomic signatures occurring in phellogen/phellem cells as compared with other tissues (e.g., Soler *et al*., 2007; Teixeira *et al*., 2018; Lopes *et al*., 2020) and developmental stages (Fernández-Piñán *et al*., 2021). Different candidate regulators of phellogen activity have been proposed, given their increased expression in developing phellem, and reported implications on meristem regulation in model plants, particularly in the vascular cambium. These include WUSCHEL-related homeobox 4 (WOX4), KNAT1/BREVIPEDICELLUS (BP), and candidate regulator AUXIN RESPONSE FACTOR 5 (ARF5), which have been recently implicated in the regulation of root phellogen activity (Xiao *et al*., 2020). These players are likely involved in an auxin-mediated pathway controlling phellogen activation, triggered by the establishment of vascular cambium in the root (Xiao *et al*., 2020). Also, members of the LATERAL ORGAN BOUNDARIES DOMAIN (LBD) family of transcription factors (TFs) that act downstream of cytokinin signaling have been shown to activate (LBD3, LBD4) and promote (LBD1, LBD3, LBD4, LBD11) secondary growth in Arabidopsis roots (Ye *et al*., 2021). Additionally, cork oak *LBD4* and *LBD1* orthologs were more expressed in root zones undergoing secondary growth, comparing to the root tip (Leal *et al*., 2022*b*). *QsLBD4* also showed increased expression in developing phellem from field-grown trees in Spring, during the period of predicted active growth (Fernández-Piñán *et al*., 2021). Cork oak genes involved in phellem differentiation have also been highlighted from transcriptomic studies, particularly regarding cell wall suberization. In addition to genes known for their role in the synthesis of suberin monomers, specific TFs such as MYB84 (QsMYB1), MYB41 and MYB93, have been validated in different systems as regulators of suberization (Kosma *et al*., 2014; Legay *et al*., 2016; Capote *et al*., 2018).

Water availability is a major determinant of cork productivity, with winter precipitation of the previous year having a strong positive impact on cork growth (Costa *et al*., 2002, 2016). However, recurrent droughts occurring in the Mediterranean regions are challenging the viability of the cork sector by reducing cork growth (and value) and affecting tree vitality (Camilo-Alves *et al*., 2017, 2020). While novel production strategies have been proposed to maximize cork oak growth (Camilo-Alves *et al*., 2020), a deeper understanding of the molecular mechanisms regulating cork development in interaction with environment is crucial to further shape more resilient plantations (e.g. through genotype selection).

Drought events are known to have severe implications on tree development. These are generally caused by hydraulic failure due to xylem cavitation, and also carbon imbalance imposed due to stomate closure and decreased CO_2_ assimilation in leaf photosynthesis (Lourenço *et al*., 2016). Regulation of the carbohydrate metabolism in adaptation to drought is an important mechanism allowing osmoregulation and refilling embolized xylem vessels (Lourenço *et al*., 2016). While this process may deplete carbon reserves in the plant, recent evidence shown that photosynthesis occurring in stems potentiates local production of carbohydrates, from CO_2_ generated by increased cell respiration taking place in stems and/or roots (Bloemen *et al*., 2013; Vandegehuchte *et al*., 2015; De Roo *et al*., 2020*a*). Depending on the species and stem diameter, the process is known to occur in green living bark tissues, and xylem (Vandegehuchte *et al*., 2015; Burrows and Connor, 2020). The suberized phellem provides an efficient barrier for CO_2_ diffusion, increasing retention of respired CO_2_ in the stems (Teskey *et al*., 2008).

In this work we conducted an experimental drought assay in cork oak saplings and targeted stems for histochemical and transcriptomic studies. We described the impact of prolonged drought on vascular cambium and phellogen activities, and identified candidate genes involved in phellem development that are downregulated by drought. We further describe and discuss the involvement of developing phellem cells in stem photosynthesis, as an integrated strategy for young cork oak plants to survive drought.

## Materials and Methods

### Plant material and growth conditions

Cork oak half-sibling acorns from two field-grown mother trees (Alcochete, Portugal, Lat. 38.747491; Long. –8.931877) were collected in November 2018 and 2020. Viable acorns were selected and cleaned by submergence in tap water, transferred to a water bath at 45 °C for 2 h and dried in a 30 °C chamber for 24 h. Acorns were stored in plastic bags at 4 °C in the dark until further use. In January 2019 (assay-1), acorns were sown on a mixture of sand and vermiculite (1:1) in 700 mL pots (1 per pot) and maintained under a 12 h photoperiod (light intensity 400 μmol m^-2^ s^-1^) at 25 °C, with regular watering. After two months, saplings were transferred to 7 L pots containing a mixture of substrate (Plantobalt) and sand (1:1) and grown from March 2019 to September 2020 in a greenhouse (Oeiras, Portugal, Lat. 38.696189, Long. –9.320762). During the first year of growth, all plants were kept under well-watered (WW) conditions (300 mL H_2_O/day) for optimal development. In March 2020, saplings were divided into two groups that were subjected to WW conditions (control) or water deficit (WD), imposed by 4 weeks of water withholding followed by 20 weeks of deficit irrigation (300 mL H_2_O/week) (Fig. 1A, Supplementary Fig. S1A). A total of 57 plants were included in each group. Soil water content (SWC) was followed during the experiment and calculated according to Sapeta *et al*. (2013). Stem diameter (1 cm above the root collar) and main stem height were measured monthly. Main stem regions developed up to 10 cm above root collar were harvested in early September 2020 (6 months after stress imposition) for histochemical and transcriptomic analysis (Supplementary Fig. S1B).

**Fig. 1.**
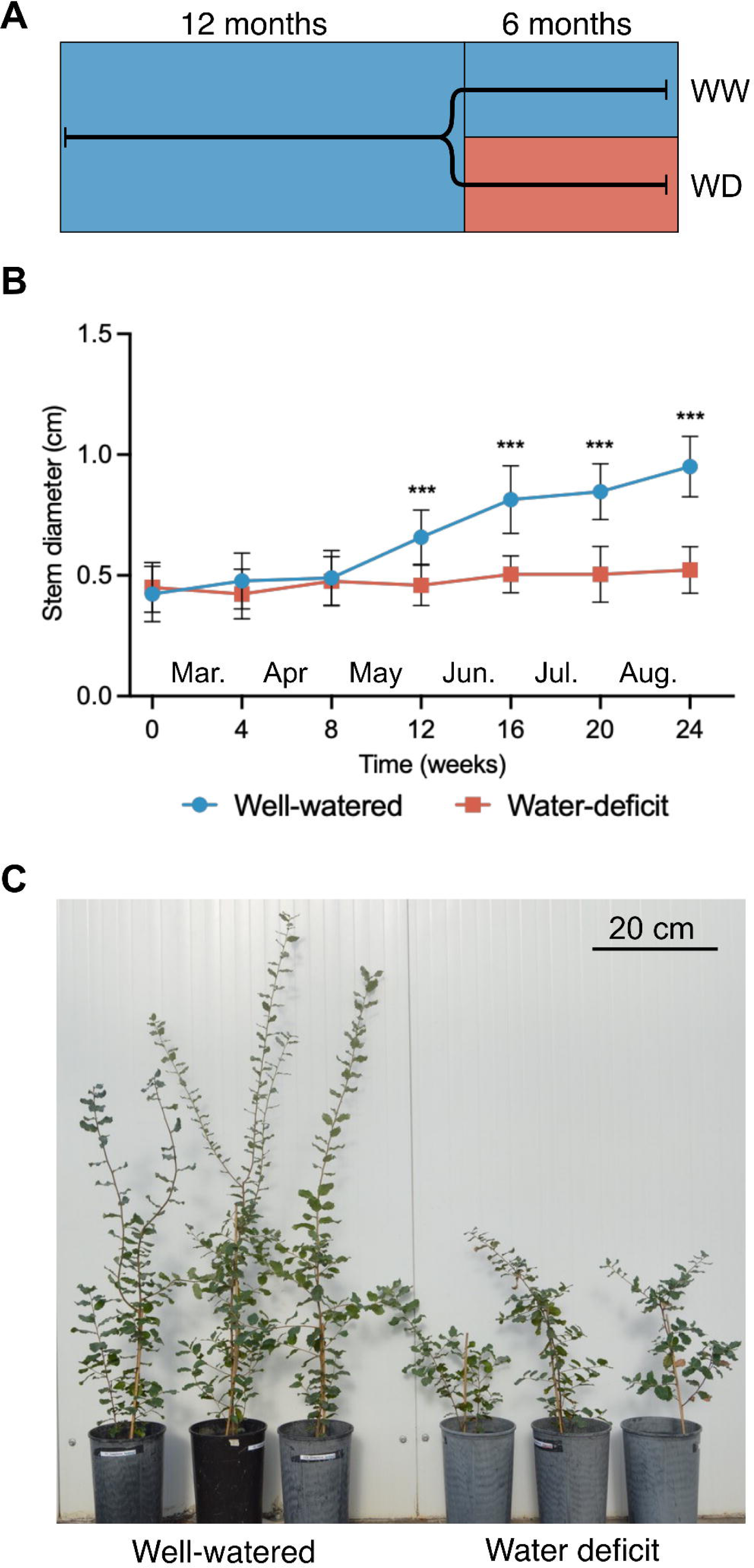
Drought assay applied to 1-year-old cork oak plants. A) Experimental design: cork oak saplings were maintained under well-watered (WW) conditions for 12 months, after which they were split in two groups, and maintained under WW or exposed to controlled water deficit (WD) conditions as described in the methods. Sampling for RNA extraction and histochemistry was performed 6 months after stress imposition. B) Stem diameter above root collar, measured monthly after stress imposition. Asterisks represent significant differences (P < 0.001, multiple t-test) between WW and WD at each timepoint. C) Representative saplings after 6 months under WW and WD conditions.

A replicate assay (assay-2) was conducted with cork oak saplings sown in January 2021. In this case, drought stress was applied as mentioned above, extending from March 2022 to August 2022. At the beginning of the assay, and to study chloroplast presence in the phellem, we prepared three saplings to be used as negative controls, by covering a 2 cm-long segment of the main stem with black tape, at exactly 2 cm above the root collar.

### Histochemical analysis and microscopy observations

For histological characterization, approximately 0.5 cm thick cross-sections of stems were sampled at 10 cm above the root collar and immediately fixed in 1% *p*-formaldehyde and 4% glutaraldehyde (in 0.1 M phosphate buffer, pH 7.2), with the application of vacuum. Stem segments were then subjected to sequential dehydration with increasing EtOH concentrations [30%, 50%, 70%, 85% and 96% (v/v)] and further embedded in Technovit 7100 resin (Heraeus Kulzer, Wehrheim, Germany) according to the manufacturer’s instructions. Semithin sections (5-10 μm thick) were obtained with a Leica RM2235 rotary microtome and fixed on a glass slide (37-40 °C for 2-3 days). For Toluidine blue (TBO) staining, cross-sections were immersed in 5% sodium hypochlorite (w/v) for 30 min, followed by immersion in 1% acetic acid (w/v) for 2 min, washed with water for 1 min, and stained with 0.05% TBO (w/v) for 1 min and washed again. After staining, samples were mounted with water and observed on a Leica DM6 B microscope under bright field. To identify suberized cells, stem cross-sections were incubated in a solution of 0.01% fluorol yellow (FY) (w/v in lactic acid) at 60 °C for 30 min, washed in water (3x 5 min each), counterstained with 0.5% aniline blue (w/v) for 30 min in the dark and washed in water for 30 min (Lux *et al*., 2005). Confocal Z-series stacks were acquired on a Zeiss LSM 880 point scanning confocal microscope using the Airyscan detector (Excitation: 480 nm; Emission: 516 nm). The Zeiss Zen 2.3 software was used to process the Airyscan raw images.

Plants from assay-2 were used for the identification of chloroplasts in phellem. The bark layer was stripped from stem segments 2 cm above the root collar, and the phellem layer (outer bark) was obtained from the natural breaking occurring at the interface with inner bark, i.e the phellogen (Supplementary Fig. S2A). Phellem layers were laid (inner side up) directly on a glass slide over a droplet of water, dried to glue in the slide, mounted with water and observed by confocal microscopy (Excitation: 458 nm; Emission: 523 nm) to detect chlorophyll autofluorescence.

Images were analysed using ImageJ software (v2.3.051). Images acquired for FY staining were used to quantify the effect of drought treatment on phellem development, by measuring the height of the phellem layer and the number of stacked suberized cells in different regions (Supplementary Fig. S2B).

### RNA extraction

For the transcriptomic analysis, xylem, inner bark (including phloem and cortex), and outer bark (phellem) layers were manually peeled from the first 10 cm-long segment of the main stems harvested above the root collar (Supplementary Fig. S1B), and frozen in liquid nitrogen. Phellem and inner bark layers were collected using the natural breaking regions occurring at the vascular cambium and phellogen, by quickly stripping the full bark layer (Supplementary Fig. S2A). Debarked stems were immediately immersed in liquid nitrogen and developing xylem was collected by scraping the wood with a scalpel blade. Each biological replicate corresponded to a pool of each tissue, collected from 3-4 stems. Frozen samples were ground with a mortar and pestle in liquid nitrogen and RNA extraction was performed according to Reid *et al*. (2006) with minor modifications described in Leal *et al*. (2022*b*). Phellem samples were ground together with 1% PVPP (w/v, considering the volume of extraction buffer). RNA quality was assessed by electrophoresis in agarose gel and quantified using a Qubit^TM^ RNA Broad Range kit and Qubit 2.0 Fluorometer (ThermoFisher Scientific, Waltham, MA, USA).

### RNA sequencing and bioinformatics analysis

RNA samples for each tissue, collected from stems grown under WW and WD were submitted to high-throughput sequencing (100 bp, paired-end reads) using DNBSEQ^TM^ at BGI Tech. (Hong Kong). Read quality was assessed using FastQC (v0.11.9). Reads were pre-processed to remove adapters and low-quality bases using Trimmomatic v0.39 (Bolger *et al*., 2014) with paired-end mode and additional parameters: LEADING:3, TRAILING:3, SLIDINGWINDOW:4:15, MINLEN:36. High-quality reads were mapped against *Q. suber* reference genome GCF_002906115.1 (CorkOak1.0) (Ramos *et al*., 2018), using STAR v2.7.7a (Dobin *et al*., 2013) with modified parameters (twopassMode = Basic, alignIntronMax = 100 000). Read counts per gene were obtained with featureCounts (Liao *et al*., 2014), using exon as feature. Differential expression analysis was conducted using the DESeq2 (Love *et al*., 2014) in R v.4.2.1. Genes with mean read count < 10 for all samples were removed. Principal component analysis was conducted using normalized counts of the top 1000 genes showing the highest variance. Differential expression analysis between WD and WW conditions and the three tissues was performed using a multifactorial design (∼Tissue + Tissue: Treatment) and the ‘apeglm’ log fold-change shrinkage estimator (Zhu *et al*., 2019). Differentially expressed genes (DEGs) found for WD *vs.* WW were filtered for adjusted *P*-value (*P*_adj_) ≤ 0.05 and fold-change (|log_2_FC| ≥ 1). Differential expression analysis between the three tissues was conducted using the Likelihood Ratio Test and DEGs were determined by adjusted *P*-value (*P*_adj_) ≤ 0.001 and baseMean > 50. DEGs between tissues were grouped using k-means clustering. Candidate *A. thaliana* orthologues were identified using Blastp analysis of cork oak predicted protein sequences on TAIR10 (Araport11).

Gene Ontology (GO) enrichment analysis of the DEGs was performed with ClueGO v2.5.9 (Bindea *et al*., 2009) plugin for Cytoscape v3.9.1, using the Arabidopsis homologs (two-sided hypergeometric test, with Bonferroni step down correction). Annotation of DEGs in specific metabolic pathways was conducted with MapMan v3.6 (Bolger *et al*., 2021), using the annotation of the cork oak predicted proteome obtained with Mercator4 (Schwacke *et al*., 2019).

### Gene expression analysis

First-strand cDNA synthesis was performed using 0.5 μg of total RNA with oligo-dT primer, using the SuperScript^TM^ III First-Strand Synthesis System (Invitrogen), following the manufacturer’s instructions. cDNA was used as the template for amplification by qPCR using gene-specific primers (listed in Supplementary Table S1). Cork oak elongation factor 1α (LOC112038967) was used as the housekeeping gene. Real-time qPCR was done in a Lightcycler 480 (Roche) using the Lightcycler 480 SYBR Green I Master mix (Roche). Target transcript abundance was calculated according to Pfaffl (2001).

### Chlorophyll fluorescence measurements

The effect of drought on photosynthesis was evaluated in assay-2 by rapid photosynthetic light curves using a MINI-PAM-II fluorometer (Heinz-Walz, Germany). This analysis was conducted in stems and leaves at day 0, 56, 100 and 120 after stress imposition. In detail, chlorophyll a fluorescence was measured in dark-adapted young leaves and stem sections (1 cm above root collar) subjected to increasing (every 20 seconds) photosynthetic active radiation (PAR, 3 to 600 μmol photons m-2 s-1).

The efficiency of Photosystem (PS) II (ΦPSII) was calculated using the formula ΦPSII = (Fm’-Fs)/Fm’ (Genty *et al*., 1989), where Fs and Fm’ represent steady state and maximum fluorescence, respectively. Electron transport rate (ETR) was calculated using ETR = ΦPSII x PAR x 0.5 x 0.84, where 0.84 is the assumed light absorbance of the leaf, and 0.5 the fraction of light absorbed by PSII (Genty et al., 1989).

## Results

### Prolonged drought inhibited secondary growth

To study the tissue-specific adaptations to drought occurring in cork oak stems, one-year-old cork oak saplings were exposed to water loss in the first 8 weeks and further maintained under approx. 10% SWC for up to 24 weeks (Fig. 1A, Supplementary Fig. S1A). Plants growing under WW conditions were maintained under 80% SWC. Under WW conditions, plants showed a constant increase in main stem height along the assay, while stem diameter started increasing after 8 weeks (Fig. 1B; Supplementary Fig. S1C). Contrastingly, WD treatment imposed a negative effect on plant development, reflected both in stem height and diameter which remained mostly unchanged along the treatment. At the end of assay-1, WW plants showed an approximately 2-fold increase in stem diameter (Fig. 1B, 2A) and height (Fig. 1C, Supplementary Fig. S1C), when compared to WD. The increase in stem diameter observed in WW plants was more pronounced in May, after the predicted seasonal reactivation of secondary growth in early Spring.

### Drought induced cell wall thickening in xylem and decreased the activity of secondary cambia

The specific impact of WD treatment in phellem development was further investigated in main stem cross-sections taken at 10 cm from root collar (Fig. 2A; Supplementary Fig. S1B), 6 months after stress imposition. At this stage we could detect clear differences in the activity of the vascular cambium and phellogen between WW and WD. Under WW, the cambial zone was clearly visible and organized, with 7-9 stacked cells (Fig. 2B), indicating increased meristematic activity. Contrastingly, the cambial zone from stems grown under WD was more compacted, with 3-4 stacked cells that were mostly flattened and smaller in volume (Fig. 2B). Looking at the phellem layer, we found less evident differences between stems under WD and WW (Fig. 2C). Yet, taking a closer look at the phellogen region we could see recent anticlinal and periclinal divisions (Fig. 2C, left panel, arrow), with the presence of non-suberized phellem cells originating from the latter. Under WD conditions, the phellogen was composed of a single cell layer located between suberized phellem and the elongated phelloderm cells (Fig. 2C, right panel, arrows). Globally, stems under WD displayed thinner periderm layers with a lower number of stacked cells, as compared to WW (Fig. 2D). Yet, this difference is modest when compared with the differences found in mean stem diameter, indicating that the negative impact of WD on secondary growth is more pronounced at the vascular cambium level than on the phellogen.

**Fig. 2.**
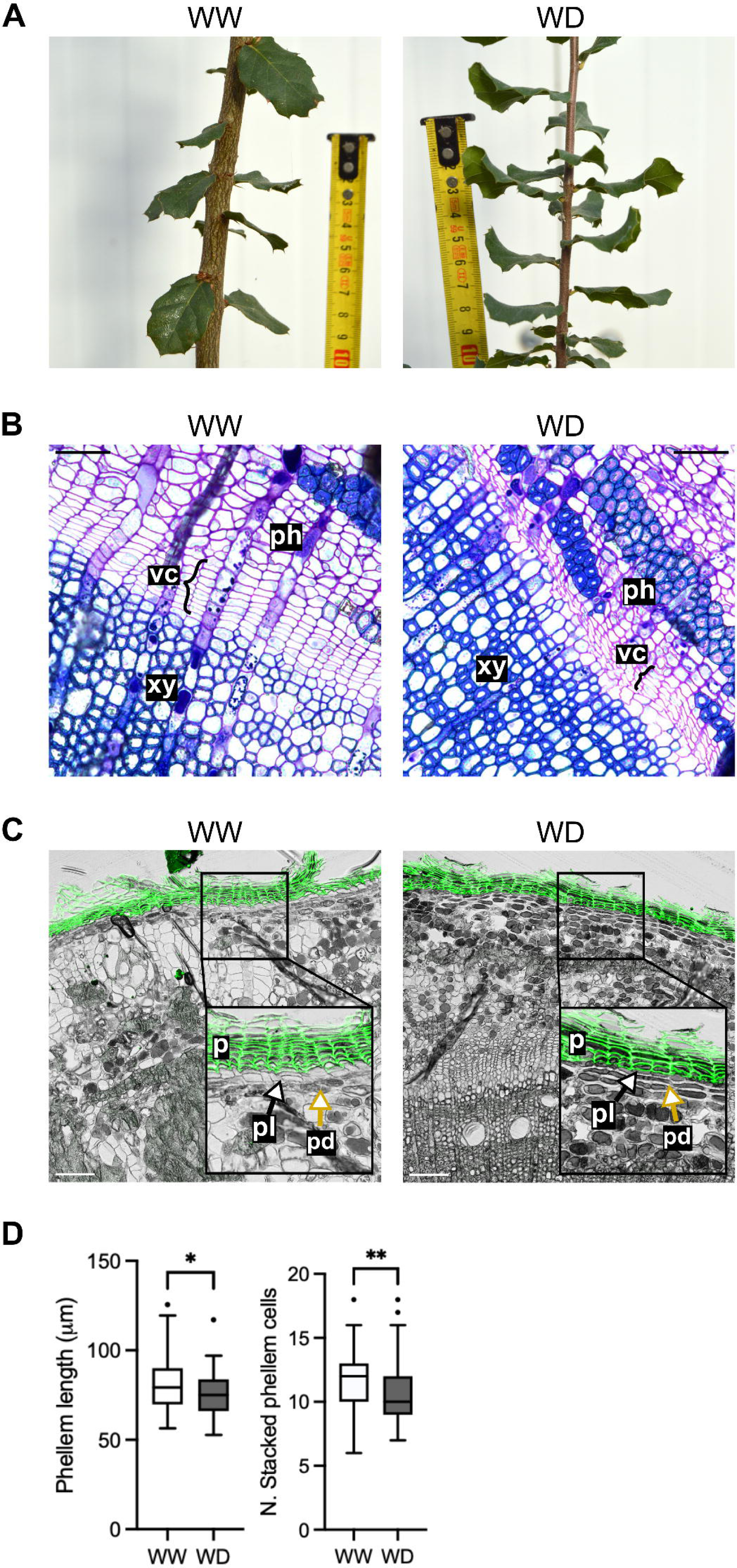
Impact of WD treatment on stem secondary growth. A) Representative pictures of the stem regions targeted for sampling, taken after 6 months under well-watered (WW) and water deficit (WD) treatments. B) Stem cross-sections stained with TBO, highlighting the vascular cambium (vc) region, developing xylem (xy) and phloem (ph); the curly brackets indicate the length of the cambial zone. Scale bars represent 50 μm. Representative pictures of 6 stems per group. C) Stem-cross sections stained with FY, highlighting the accumulation of suberized cells in phellem (p), derived from phellogen (pl) activity. Inset images are details of the indicated areas, also highlighting the location of phelloderm (pd). Scale bars represent 100 μm. Representative pictures of 6 stems per group. D) Quantification of phellem length (left) and number of stacked phellem cells (right) in 6 stems per group. For each image, phellem length and number of cells were measured from 4 different positions, and boxplots represent the distribution of all the measurements per group. Asterisks represent significant differences (* P < 0.05, ** P < 0.01, Mann-Whitney test) between WW and WD.

### Tissue-specific transcriptomic dynamics highlight candidate genes involved in phellogen activity

To get further insights into the genetic players involved in drought adaptation in stems, we performed a genome-wide transcriptomic analysis following a tissue-specific approach. Main stems collected up to 10 cm above root collar (Supplementary Fig. S1B) were manually peeled, using the natural breaking points occurring at the vascular cambium and the phellogen, to obtain three main layers for RNA extraction – xylem, inner bark, and phellem (outer bark). Since the phellogen is commonly a single layer of cells, we sought to determine if these cells would stay attached to the inner bark, by analysing stem cross-sections with and without phellem (Supplementary Fig. S2C). In cross-sections without phellem we could detect partial removal of phellogen cells, with some regions showing no traces of this cell type (Supplementary Fig. S2C, red arrows), indicating that these meristematic cells were collected attached to the phellem.

RNA-seq analysis yielded an average of 25.5 M read pairs per library, of which 18.5 M could be uniquely mapped and assigned to genes (Supplementary Table S2). Principal component analysis based on the transcriptomic profiles clearly separated phellem samples from those collected from xylem and inner bark along PC1 (Fig. 3A). Variability observed between WW and WD conditions was mostly represented along PC2.

**Fig. 3.**
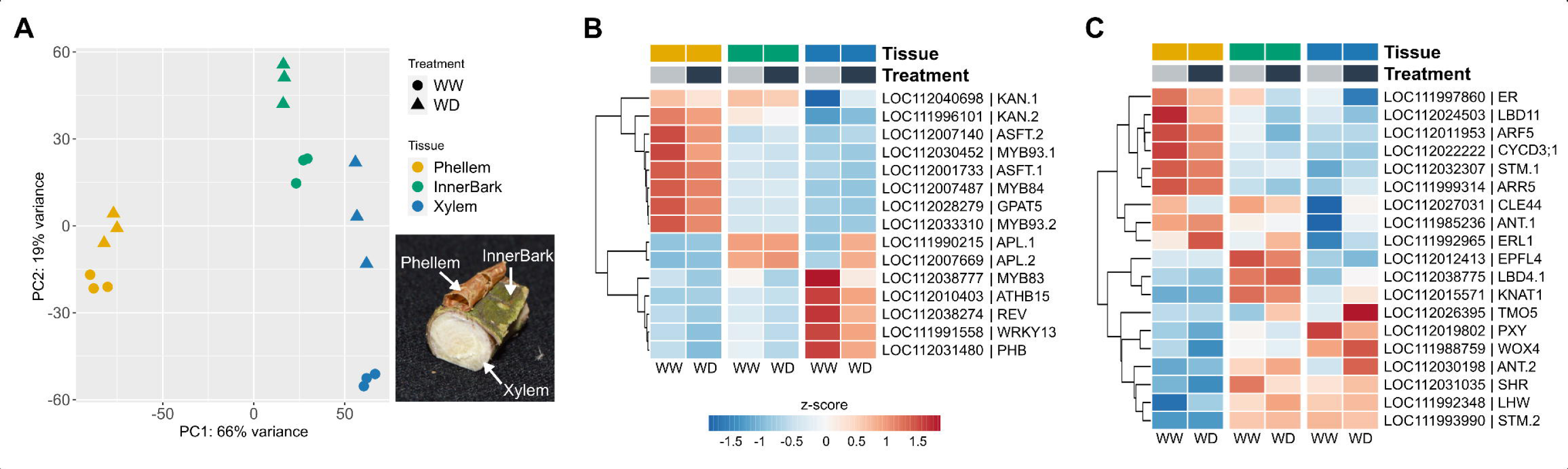
Tissue-specific transcriptional dynamics occurring in phellem, inner bark and xylem from young cork oak stems developed under well-watered (WW) and water deficit (WD) conditions. A) Principal component analysis of the stem samples targeted for RNA-seq, including three biological replicates per tissue and condition. B) Heatmap including candidate genes involved in phellem and vascular tissue differentiation that are differentially expressed between tissues. C) Heatmap including candidate genes involved in meristem activity, that are differentially expressed between tissues (Likelihood Ratio Test, adjusted p-value < 0.001). Gene expression is represented by normalized read counts (vst) and centred to the mean expression per gene (z-score).

To further confirm tissue specificity and assess the putative representation of the vascular cambium and phellogen we explored the diversity of differentially expressed genes between the three layers, independently of the growth conditions. We identified 8550 DEGs between at least one of the three possible pair-wise comparisons (phellem *vs.* inner bark, phellem *vs.* xylem, and inner bark *vs.* xylem), which were further grouped in six clusters based on mean normalized expression (Supplementary Fig. S3, Supplementary Table S3). Among the genes with contrasting expression profiles between tissues we found multiple genes with predicted roles in cellular differentiation and regulation of meristematic activity (Fig. 3B, C). As expected, the expression of genes involved in suberin synthesis was enriched in phellem, including *QsGPAT5* (LOC112028279), *QsASFT.1/2* (LOC112001733, LOC112007140)*, QsMYB93.1/2* (LOC112030452, LOC112033310), and *QsMYB84* (LOC112007487), which is a homolog of *QsMYB1* (57.26% amino acid identity in Blastp). In turn, cork oak homologs of TFs with known roles in the differentiation of xylem -*QsMYB83* (LOC112038777), *QsATHB15/CORONA* (LOC112010403), *QsREVOLUTA* (LOC112038274), *QsPHABOLOSA* (LOC112031480) and *QsWRKY13* (LOC111991558) - and phloem - *ALTERED PHLOEM DEVELOPMENT* (*QsAPL.1/2,* LOC111990215, LOC112007669) and *KANADI* (*QsKAN*) - were more active in these specific layers. Interestingly, the two paralogs of KANADI, *QsKAN.1/2* (LOC111996101, LOC112040698) were also expressed in the phellem layer (Fig. 3B).

Among the DEGs found between the three layers we also found different genes with predicted roles in meristem development. Expression of cork oak homologs of the *QsPXY* (LOC112019802) and *QsWOX4* (LOC111988759), which are regulators of cambial cell proliferation were detected in the xylem layer, while a CLAVATA3/ESR (CLE)-related protein 44 (*QsCLE44,* LOC112027031) homolog was enriched in the inner bark (Fig. 3C), in agreement with the phloem-specific expression determined in model plants. Other genes with predicted roles in the control of the vascular cambium, such as *QsLBD11* (LOC112024503), *QsARF5* (LOC112011953), *QsARR5* (LOC111999314) and the cyclin-dependent kinase *QsCYCD3;1* (LOC112022222) showed enriched expression in the phellem layer (cluster 3 and 6, Fig. S3, Supplementary Table S3). This evidence indicates that the phellem and xylem layers are likely also representing the transcriptome of the corresponding cambial tissues.

### Drought negatively impacts the expression of candidate regulators of phellogen activity

The transcriptomic profile of samples collected under WD conditions was compared to that of WW to assess the impact of drought on tissue-specific pathways. In the phellem layer we found 1481 differentially expressed genes (DEGs) between WD and WW (Supplementary Table S4), of which 606 were upregulated and 875 were downregulated. In the inner bark and xylem we found 2062 (561 up- and 1501 downregulated) and 5275 (2593 up- and 2682 downregulated) DEGs (Supplementary Tables S5-S6). These results indicate a major impact of drought conditions in xylem when compared to the bark layers. From the DEGs determined in phellem, we selected 9 DEGs for validation by real-time qPCR. From these, 6 were also differentially expressed in inner bark and xylem. Comparison of the log2FCs ratios (WD *vs.* WW) determined by RNA-seq and qPCR for the three tissues revealed similar expression trends, supporting the RNA-seq results (Supplementary Fig. S4A).

Gene ontology (GO) enrichment analysis showed a clear representation of genes acting in cell wall biogenesis, particularly in glucan and pectin metabolism, and phenylpropanoid biosynthesis in DEGs from all three tissues (Fig. 4, Supplementary Tables S7-S9). Other DEGs annotated in these two categories were also identified more specifically in xylem and inner bark. DEGs acting in fatty acid biosynthesis were enriched in inner bark and phellem (Fig. 4). DEGs involved in cell wall biogenesis, and phenylpropanoid/fatty acid biosynthesis were mostly downregulated under WD (Supplementary Fig. S4B-D, Fig. 5A). DEGs with predicted roles in meristem/tissue development were mostly represented in the xylem, while genes involved in cell cycle processes were enriched in DEG groups determined for xylem and inner bark (Fig. 4). DEGs included in these functional categories were also mostly downregulated (Supplementary Fig. S4C,D). Interestingly, DEGs found in phellem, showed an enrichment in photosynthesis-related genes which were mostly upregulated under WD (Supplementary Fig. S4B), with some of them shared with inner bark (Fig. 4). Genes involved in response to abiotic stress (including ABA and osmotic stress) were also enriched in DEGs found in all three tissues (Supplementary Fig. S4B-D), although these categories were more represented in xylem (Fig. 4).

**Fig. 4.**
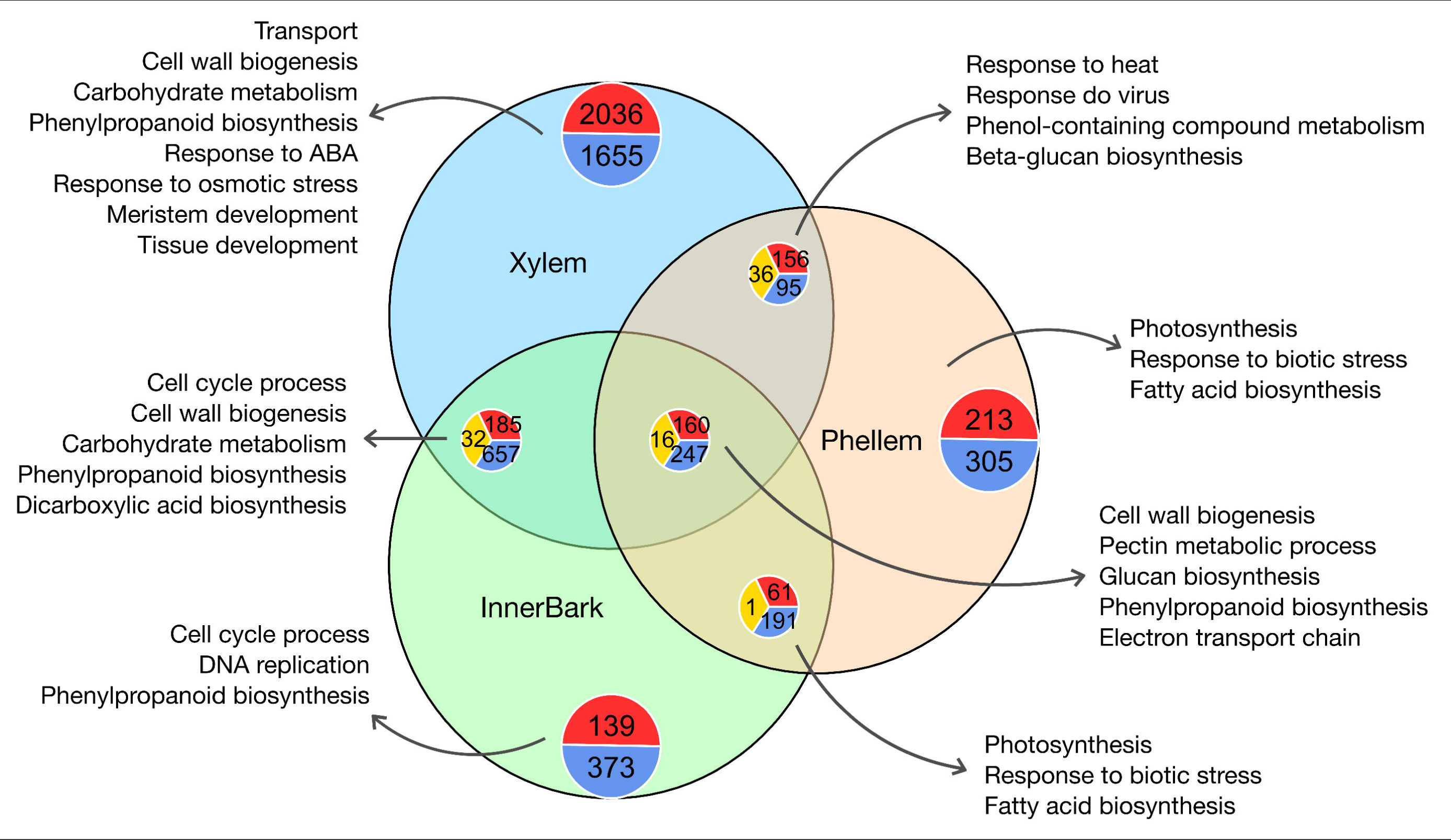
Common and tissue-specific changes occurring in cork oak stems in adaptation to WD. The Venn diagram was built using the list of DEGs for WD *vs.* WW conditions, determined for xylem (blue), inner bark (green) and phellem (yellow). The pie charts represent the number of genes upregulated (red), downregulated (blue) or showing contrasting patterns between tissues (yellow).

**Fig. 5.**
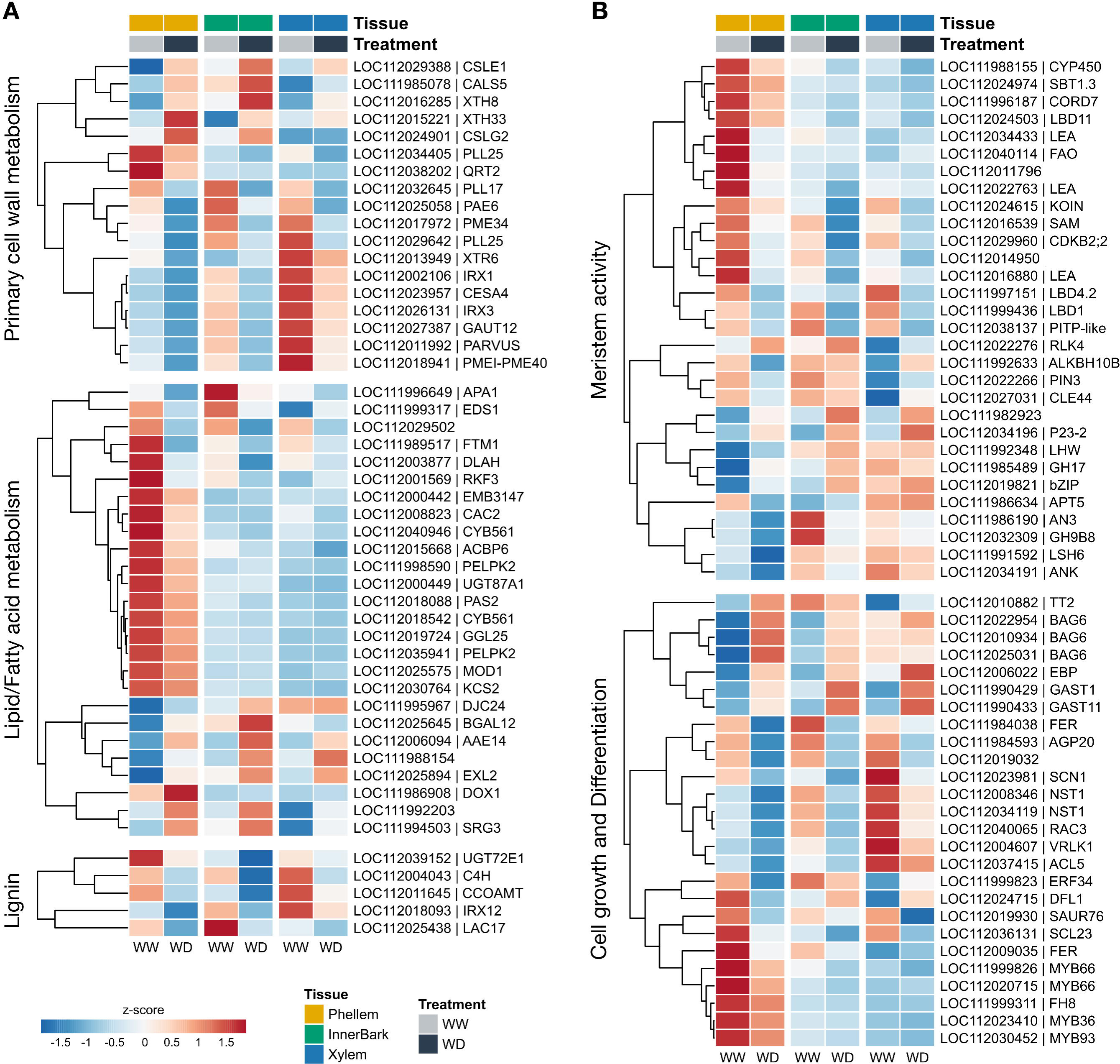
Heatmap representing the expression profile of selected DEGs determined for phellem under WD *vs.* WW, organized by their predicted role in (A) cell wall biogenesis and (B) meristem regulation and cell division. The expression of these genes in inner bark and xylem is also displayed to highlight tissue-specific trends. The heatmap was built using the mean normalized expression of the biological replicates, and centred to the gene mean expression in all samples (z-score). Additional genes for similar categories are represented in Supplementary Fig. S5.

Looking in more detail at DEGs found in phellem and acting during cell wall establishment, we have identified the upregulation of two cellulose synthase-like genes *QsXTH8* and *QsXTH33* (LOC112029388 and LOC112024901), two xyloglucan endotransglucosylase/hydrolase genes *QsCSLE1* and *QsCSLG2* (LOC112016285, LOC112015221) involved in hemicellulose synthesis, and one callose synthase (LOC111985078) (Fig. 5A). Other genes with predicted roles in primary cell wall biogenesis or modification were downregulated in phellem although most showed an overall higher expression in xylem and inner bark layers. The exceptions were *QsXTH33*, *QsCSLG2*, and two downregulated genes involved in pectin metabolism, polygalacturonase (LOC112038202) and one paralog of pectate lyase PLL25 (LOC112034405) (Fig. 5A). DEGs involved in lipid/fatty acid metabolism were enriched in phellem, with a majority being downregulated under WD and showing reduced expression in the xylem and inner bark (Fig. 5A; clusters 1, 3 and 6, Fig. S3, Supplementary Table S3). DEGs acting in lignin metabolism were also found in phellem, including a UDP-glucosyl transferase (LOC112039152) that also showed enriched expression in phellem (Fig. 5A), as compared to the other layers (cluster 3, Fig. S3, Supplementary Table S3). This is also the case of other DEGs with predicted roles in cell wall biogenesis or organization, such as the transmembrane ascorbate ferrireductase *QsACYB-2* (LOC112005119), the heavy-metal associated protein *QsHMP24* (LOC112035134), and a hypothetical protein *LOC112023550* (Fig. S5A, Fig. S3, Supplementary Table S3).

The morphological analysis of outer bark from stems maintained under WW and WD suggested a shut-down of phellogen activity. Therefore, we searched for candidate regulators of phellem formation based on functional annotation. Among the DEGs determined in the phellem between WD and WW, we found 30 genes with predicted roles in the regulation of meristem activity, which were mostly downregulated under WD (Fig. 5B). This was the case of *QsLBD1* (LOC111999436), *QsLBD4.2* (LOC111997151) and *QsLBD11*, which also showed different patterns of expression across the stem layers. While the *QsLBD1* was downregulated in all the three layers, *QsLBD4* was only downregulated in phellem and xylem, while *QsLBD11* was mostly expressed in phellem (Fig. 5B). Expression of *QsCLE44*, *QsAPT5* (LOC111986634), and *QsPIN3* (LOC112022266) was downregulated only in phellem, in addition to two *LEA* protein-coding genes (LOC112034433, LOC112022763), *CORTICAL MICROTUBULE DISORDERING7* (*QsCORD7,* LOC111996187), *FAD-dependent oxidoreductase* (*QsFAO*, LOC112034433), and *subtilase QsSBT1.3* (LOC112024974). Regarding genes associated with cell growth and differentiation, we found the phellem-specific downregulation of candidate regulators of cell wall suberization, namely *MYB93* and *MYB36* (LOC112023410) homologs, in addition to two *WEREWOLF* paralogs involved in cell fate determination (*QsMYB66.1/2,* LOC111999826 and LOC112020715) and a formin-like protein (*QsFH8*, LOC111999311) involved in actin cytoskeleton organization. Meristem-related genes that were upregulated in phellem under WD included *RECEPTOR-LIKE PROTEIN KINASE 4* (*QsRLK4,* LOC112022276), co-chaperone *QsP23-2* (LOC112034196), *O-Glycosyl hydrolase 17* (*QsGH17*, LOC111985489), and a *bZIP* TF (LOC112019821), which were also upregulated in the xylem and/or inner bark (Fig. 5B), and may be involved in growth inactivation. We also found a strong upregulation of three *Bcl-2-associated athanogene 6* chaperone gene paralogs (*QsBAG6*), *TRANSPARENT TESTA2* (*QsTT2*, LOC112010882), ethylene-responsive factor *QsEBP* (LOC112006022), two gibberellin-related *QsGAST1* (LOC111990429) and *QsGAST11* (LOC111990433), and *ACT domain repeat 1* (*QsACR1*, LOC112028560) (Fig. 5B, Supplementary Fig. 5B). *QsGAST11* was one of the genes showing the highest fold-change in all three tissues (Supplementary Tables S4-S6). As observed with *QsGAST11*, *QsGAST1* and *QsEBP* were also upregulated in the three tissues (Fig. 5B). In turn, *QsTT2* was upregulated in phellem and xylem, *QsBAG6* genes were upregulated in phellem and inner bark, and *QsACR1* was only upregulated in phellem (Supplementary Fig. S5B).

The transcriptomic data agrees with the morphological study performed in stems grown under WW and WD, highlighting a negative impact on the expression of genes associated with stem development and secondary growth. This includes regulation of cambial activity, cell growth and differentiation, matching with a reduction in the expression of genes involved in cell wall biogenesis. We also identified multiple candidate genes that may play a role in the negative regulation of growth and/adaptation to drought conditions.

### The phellem layer participates in drought-responsive bark photosynthesis

The RNA-seq analysis conducted in phellem highlighted an enrichment in photosynthesis-related DEGs upregulated under drought (Fig. 4). Woody/bark-tissue photosynthesis is already documented in several tree species, as a way to recycle respired CO_2_ in the stem and mitigate carbon starvation under drought conditions (Vandegehuchte *et al*., 2015). Yet, this is empirically assumed to occur in green tissues from the inner bark of young stems with thin outer bark. In agreement, most of the photosynthesis related DEGs found in phellem were also induced and highly expressed under WD conditions in cork oak inner bark (Supplementary Fig. S5C). These genes are included in different pathways associated with photosynthesis, namely the Calvin cycle, chlororespiration, photosystem I and II components and light-harvesting complexes (Fig. 6). Interestingly, genes associated with NDH-dependent cyclic electron flow (CEF) were also upregulated (*QsNDF4*; *QsNDHL*; *QsNdhU*, *QsNDF5*; Supplementary Fig. S5C; Supplementary Tables S4 and S5). Based on these results we hypothesized that developing phellem cells could participate in bark photosynthesis.

**Fig. 6.**
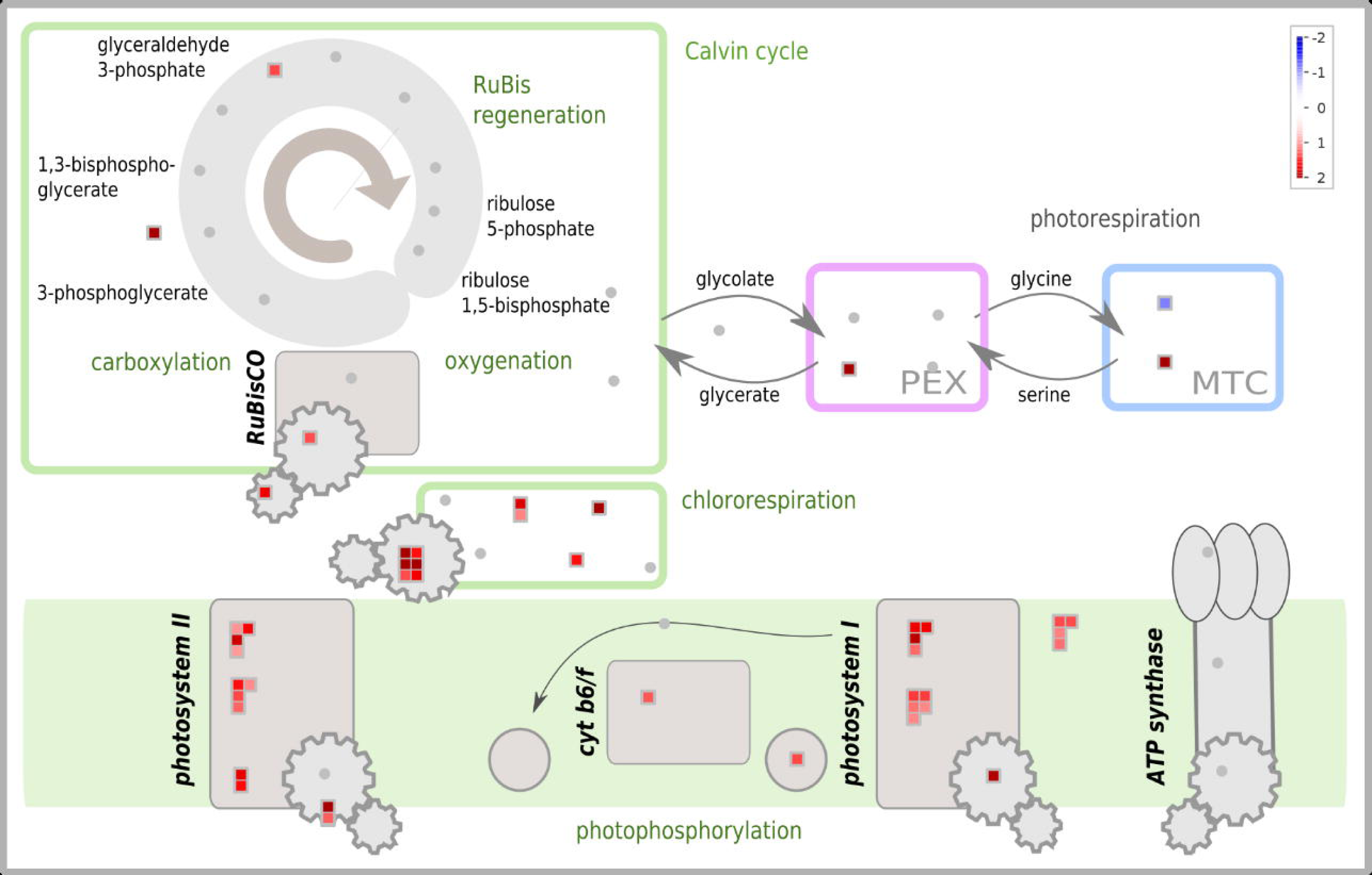
MapMan diagram representing photosynthesis-associated DEGs found in phellem in response to WD conditions (according to functional annotation using Mercator 4). Colours represent the log2FC determined in phellem for the annotated genes.

To validate the activation of bark photosynthesis in response to drought, we repeated the drought experiment (assay-2) and assessed photosynthetic activity in leaves and main stems (measured above the root collar) at different stages. In leaves, photosynthesis efficiency under WD remained lower than that of WW conditions along the three stages characterized (Fig. 7A, Supplementary Fig. S6). In stems, photosynthesis efficiency under WW was globally lower than the one detected in leaves (Supplementary Fig. S6). At 56 days of WD treatment, a decrease in photosynthesis efficiency was detected when compared to control, in agreement with the results obtained for leaves (Fig. 7A). Yet, at later stages (D100 and D120) photosynthesis efficiencies in stems under WD and WW were similar, suggesting a compensatory effect in adaption to drought. This agrees with the increase in photosynthesis related genes detected by RNA-seq.

**Fig. 7.**
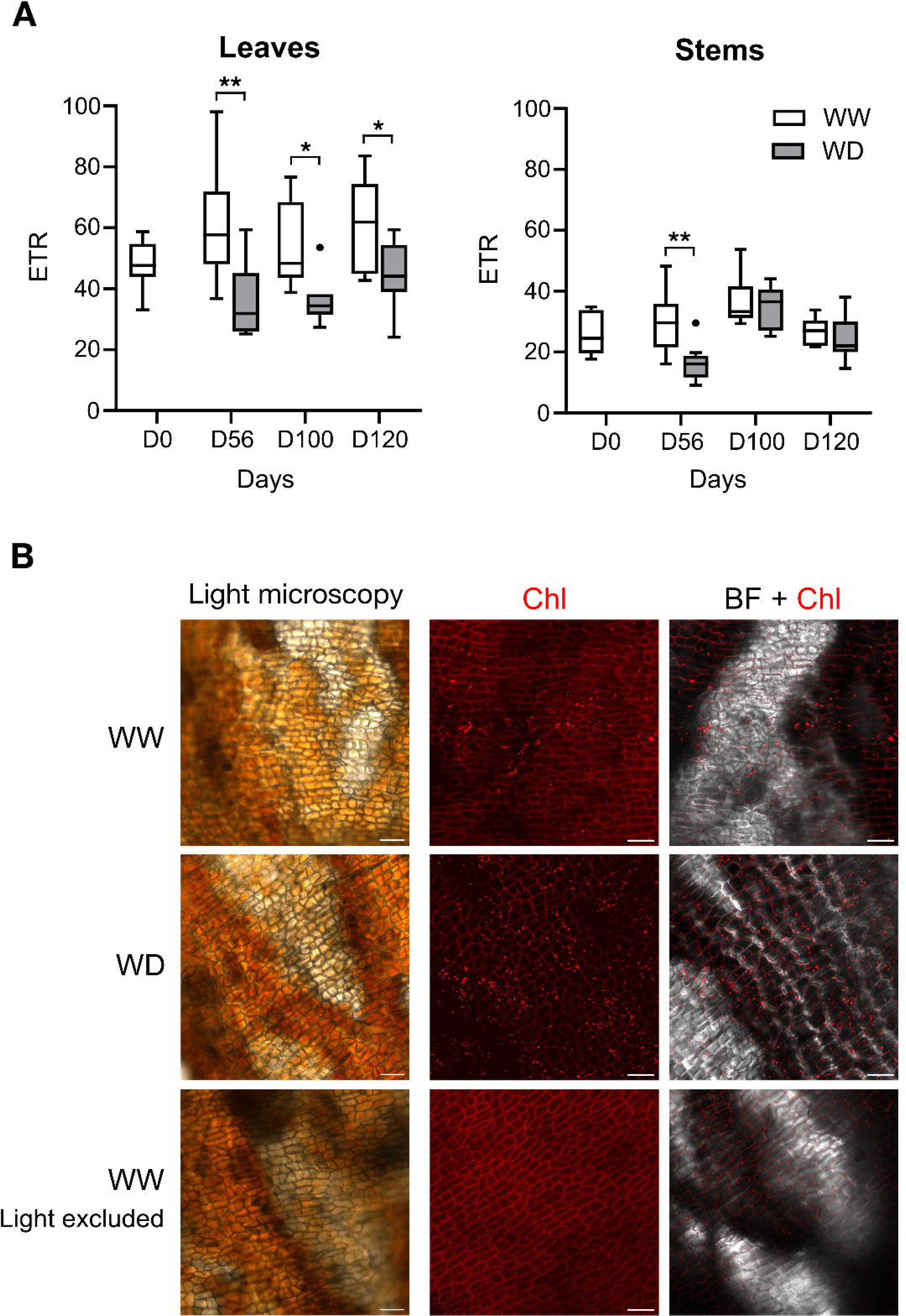
Phellem cells play a role in bark photosynthesis. (A) Mean ETR (μmol electrons m^-2^ s^-1^), measured at maximum irradiance (601 ± 59.13 μmol photons m^−2^ s^−1^) in leaves and main stems (above root collar), during different stages of WD treatment (Day 0, 56, 100 and 120), in comparison to WW conditions. Asterisks indicate statistical significance for the comparison of means between WD and WW groups, n = 6-13 plants (* P < 0.05, ** P < 0.01, t-test). (B) Chlorophyll fluorescence observed at the inner side of phellem layers stripped from the main 1.5 year-old stems. The left pane shows representative images of the phellem layer in light microscopy observations. In the right pane, the first column shows representative confocal microscopy images of chlorophyll fluorescence (Chl) in the phellem layer (Excitation: 458nm; Emission: 523nm) and the second column shows the corresponding images merged with the bright field (BF). Additional replicates are included in Supplementary Fig. S7. The light-excluded row refers to the phellem layer collected from a main stem region (above the root collar) that was covered for 120 days with black tape (Fig. S7A) – only autofluorescence from cell walls was detected in this sample. Scale bar: 50 μm.

At D120, we manually removed the phellem layer from stems and analysed them under a confocal microscope to look for the presence of chloroplasts. As a negative control, we analysed the phellem layer from a stem segment (WW condition) that was covered with black tape for 120 days (Supplementary Fig. S7A). By analysing chlorophyll fluorescence, we detected the presence of chloroplasts in phellem samples from WW and WD (Fig. 7B). In turn, no chlorophyll autofluorescence was detected in the light-excluded stem sample. Additionally, we found that the number of chloroplasts was higher in WD when compared to WW samples (Fig. 7B; Supplementary Fig. S7B). Our results support that developing phellem is photosynthetically active and participates in the adaptive response to drought by modulating photosynthesis.

## Discussion

Controlled irrigation or fertigation in cork oak stands has been proposed as a viable solution to improve cork oak development and decrease to half the time of first harvest (Vessella *et al*., 2010; Camilo-Alves *et al*., 2020, 2022). This time is usually determined by tree stem circumference at breast height (>70cm), which under natural (and optimal) conditions usually occurs after 20 years’ age or more (Pereira, 2007). Fertigation campaigns applied from late Spring to Autumn in adult trees contributed to both cork and wood radial growth increments (Camilo-Alves *et al*., 2022; Poeiras *et al*., 2022).

In the present work we conducted an experimental drought assay using young cork oak plants, to characterize the impact on stem secondary growth at cellular and molecular levels. As expected, the prolonged drought treatment had negative consequences on plant development, limiting both primary and secondary growth. Under WW conditions, the increase in stem diameter was observed after May, marking the seasonal reactivation of secondary growth. At this stage, plants under WD treatment were already adjusted to minimum SWC, which correlated with the inhibition of plant growth (Fig. 1, Supplementary Fig. S1). In fact, in WD mean stem diameter was maintained during the 6 months. Through histochemical analysis, we showed that the negative impact of WD on radial stem growth was more pronounced on the vascular cambium than on the phellogen (Fig. 2). The vascular cambium is known to regulate phellogen in Arabidopsis roots, since mutants with impaired vascular cambium activity indirectly affect phellogen activity (Xiao *et al*., 2020). This may also be the case when plants are adapting to drought. Also, we hypothesize that under WW conditions, the phellogen might have invested in anticlinal divisions to cope with the rapid increase in radial growth. This scenario might explain the modest increase in phellem thickness observed under WW conditions, in the 4-month time frame of observed active secondary growth.

By applying a tissue-specific approach and comparing samples under active and inactive growth, our transcriptomic study allowed assessing common and specific gene expression trends along the three layers of the stem. By analyzing the expression pattern of different marker genes, we confirmed the specificity conferred by the manual peeling of the different layers (Fig. 3). In inner bark (containing the phloem) and xylem layers, we confirmed the differential expression of the *CLE44/PXY/WOX4* module, described to regulate vascular cambium differentiation (Ruonala *et al*., 2017; Turley and Etchells, 2021). PXY and WOX4 are marker genes for the vascular cambium in Arabidopsis (Suer *et al*., 2011; Xiao *et al*., 2020) and the high expression of *QsPXY* and *QsWOX4* in the xylem was indicative of an enrichment of cambial cells in this layer. CLE44 encodes a signaling peptide that travels from the phloem to interact with PXY and regulate cambial activity (Ruonala *et al*., 2017; Turley and Etchells, 2021). *CLE44* promoter activity was detected in phloem, but also in pericycle of Arabidopsis roots (Hirakawa *et al*., 2010), which is the precursor of phellogen in this organ (Wunderling *et al*., 2018; Leal *et al*., 2022*a*). In cork oak stems, *QsCLE44* was enriched in inner bark layer, but also in the phellem, in comparison to xylem (Fig. 3C). This pattern was also found between phellem and xylem layers in adult cork oak trees (Lopes *et al*., 2020), supporting a the hypothesis of regulatory role of this gene in phellem development.

The *QsLBD11*, *QsARF5*, *QsARR5* and *QsCYCD3;1* showed enriched expression in the phellem layer when compared to the other layers (Fig. 3C). ARF5 is an auxin-dependent regulator of phellogen activity in Arabidopsis roots (Xiao *et al*., 2020). *CYCD3;1* and *ARR5* are cytokinin-responsive genes previously implicated in the regulation of secondary growth and were validated as markers for phellogen in Arabidopsis roots and potato tubers (Randall *et al*., 2015; Vulavala *et al*., 2019). LBD11 was recently implicated in a feedback loop involving reactive oxygen species (ROS) that regulate vascular cambium redox homeostasis and radial growth in Arabidopsis. Yet, in this species, LBD11 promoter activity was mostly prevalent in vascular cambium and xylem parenchyma of the hypocotyl (Dang *et al*., 2023). Contrastingly, the increased expression of *QsLBD11* in phellem layer, as opposed to vascular tissues, was also detected in adult cork oak trees (Lopes *et al*., 2020). Based on the downregulation of *QsLBD11* under WD (Fig. 5B), we hypothesize that this gene could be involved in the regulation of phellogen activity.

Another interesting aspect highlighted by this study is the putative functional diversification of cork oak paralogs of *SHOOT MERISTEMLESS* (*STM*), *AINTEGUMENTA* (*ANT*) and *LBD4* TFs, which are also regulators of cambial activity (Liebsch *et al*., 2014; Randall *et al*., 2015; Ye *et al*., 2021). Expression of *QsSTM.1* (LOC112032307) and *QsANT.1* (LOC111985236) was higher in phellem layer, while *QsSTM.2* (LOC111993990) and *QsANT.2* (LOC112030198) was more prevalent in inner bark and xylem (Fig. 3C). In turn, *QsLBD4.1* (LOC112038775) expression was higher in the inner bark layer (Fig. 3C), while *QsLBD4.2* (LOC111997151) was more expressed in phellem and xylem layers under WW (Fig. 5B), being downregulated by WD. Functional diversification of paralog genes has been already reported between annual and perennial plants as a result of genome duplication events. One pivotal example is the duplication of FLOWERING LOCUS T (FT) in poplar, with FT1 regulating meristem transition to reproductive stage and FT2 controlling vegetative growth (Hsu *et al*., 2011). Further functional and evolutionary studies will be important to validate the impact of duplication events in the adaptation of secondary growth to the perennial habit.

In agreement with the evident growth arrest observed in adaptation to drought, all stem layers showed a downregulation of genes involved in the growth and differentiation of vascular tissues and phellem. These mostly included genes involved in cell wall biogenesis, from primary to secondary cell wall components. More specifically, cork oak homologs of the regulator of suberization *MYB93*, and genes acting in the synthesis of aliphatic and aromatic components of suberin were downregulated by WD (Fig. 5B). Other regulators of cell differentiation downregulated by WD included cork oak homologs of SCARECROW-LIKE23 (LOC112036131) and MYB36, which are involved in root endodermis differentiation (Kamiya *et al*., 2015; Long *et al*., 2015), and MYB66 that acts on epidermal cell fate determination (Kwak *et al*., 2015). As cork oak productivity is mainly related to phellem annual growth, we also looked for candidate genes involved in meristem activity and regulation of growth. Given their downregulation under WD we propose *QsLBD1*, *QsLBD4.2*, *QsLBD11*, *QsCLE44* and the predicted auxin transporter *QsPIN3* as positive regulators of phellem growth. Interestingly, Arabidopsis *PIN3* is downregulated in roots in response do osmotic stress, leading to growth inhibition (Yuan *et al*., 2021).

Contrastingly, among the genes upregulated under WD we found two gibberellin-responsive genes, *QsGAST1* and *QsGAST11*, which share high homology to the plant antimicrobial peptide Snakin2 from potato (53.5% and 76.2% aa identity, respectively). StSnakin2 is negative regulator of H_2_O_2_ accumulation and biosynthesis of lignin precursors in the tuber periderm, also playing a determinant role in inhibiting tuber sprouting (Deng *et al*., 2021). Additionally, three *QsBAG6* paralogs were strongly expressed in cork oak stems and also upregulated under WD in the phellem and inner bark. The BAG gene family encode highly conserved molecular chaperone cofactors that have been implicated in plant growth, autophagy, and response to (a)biotic stresses (Jiang *et al*., 2023). BAG6 proteins include a Calmodulin-binding motif and have been implicated in autophagy in response to fungi, but also flowering and response to heat stress (Jiang *et al*., 2023). These genes are likely involved in an adaptation mechanism to WD conditions that may also negatively control growth.

Additionally, a strong up-regulation of genes related to photosynthesis was found under WD in the inner bark and phellem, including PSII and PSI components and respective light-harvesting systems (Fig. 6). In agreement, while photosynthetic activity in stems decreased at early stages of adaptation to WD, we detected an adjustment at later stages that reached the levels measured under WW conditions (Fig. 7A). These results suggested an increase in light harvesting and photosynthetic capacity of stems in adaptation to long-term exposure to WD. Refixation of internal CO_2_ through photosynthesis in the green tissues of the stems is particularly important under water deficit conditions, by filling local carbon demands necessary for plant metabolism, allowing turgor maintenance and osmoprotection, and thus contributing to hydraulic integrity (De Roo *et al*., 2020*a*,*b*; Rodríguez-Calcerrada *et al*., 2021; Trifilò *et al*., 2021). Since stem photosynthesis is frequently associated with green bark issues, the upregulation of photosynthesis-related genes in phellem prompted us to look at chlorophyll fluorescence in this layer. Surprisingly, we detected the presence of chloroplasts in developing phellem cells, with increased abundance in WD conditions (Fig. 7B), indicating an active participation of phellem in bark photosynthesis.

An up-regulation of genes associated with the NDH-dependent CEF around PSI was also observed in the two bark tissues. Increased CEF was reported in the branches of cotton in response to high light, protecting the photosynthetic apparatus by the increase of heat dissipation under stress (Li *et al*., 2023). Similar observations have been made for leaves of several species in response to drought (Suorsa, 2015). The increased expression of these CEF-associated genes suggests that CEF increases in young cork oak stems in response to WD. Similarly to what is described for leaves, increased CEF would allow a better adjustment between CO_2_ fixation and photoprotection. Yet, considering the low light incidence in the lower part of the stem that was sampled in our study, we hypothesize that CEF is increased to allow higher ATP production that may be needed for adjusting the ATP/NADH demand of the downstream metabolism (Huang *et al*., 2015).

Cork oak is described as a isohydric species, tightly regulating stomatal conductance to prevent xylem cavitation, being more susceptible to carbon starvation (Camilo-Alves *et al*., 2017). Altogether, our results highlight the dynamic role of bark photosynthesis in young cork oak trees to withstand water deficit during the establishment years. We showed that the phellem is photosynthetically active and participates in the adaptive response to drought. Considering the expected exacerbation of drought and heat events due to climate alterations and associated implications in tree vitality, the carbon gain resulting from bark photosynthesis may be crucial to avoid young tree mortality. The identification and selection of resilient genotypes with enhanced bark photosynthesis could play a pivotal role in mitigating the decline of cork oak forests.

## Abbreviations

CE: cyclic electron flow
ETR: electron transport rate
DEGs: differentially expressed genes
FY: fluorol yellow
GO: gene ontology
SWC: soil water content
TBO: toluidine blue
TFs: transcription factors
WW: well-watered
WD: water-deficit.

## Supplementary data

The following supplementary data are available at JXB online:

**Fig. S1.** Soil water content and cork oak stem height measured along the drought treatment (assay-1).

**Fig. S2.** Isolation and measurement of the phellem layer in young cork oak plants.

**Fig. S3.** K-means clustering of DEGs found between tissues, independently of the treatment.

**Fig. S4.** Validation of DEGs by qPCR, Gene Ontology and Mapman functional annotation and enrichment analysis.

**Fig. S5.** Heatmaps representing the expression profile of selected DEGs determined for phellem under WD *vs.* WW.

**Fig. S6.** Electron transport rate rapid light curve response measured in leaves and stems along the drought treatment (assay-2).

**Fig. S7** Chlorophyll fluorescence observed at the inner side of phellem layers stripped from main 1.5 year-old stems.

**Table S1.** Accession numbers and primer sequences of the target DEGs used for qPCR analysis.

**Table S2.** Number of raw and high-quality reads generated by Illumina sequencing and mapping efficiencies.

**Table S3.** List of DEGs calculated between the 3 tissues (phellem, inner bark and xylem) independently of the growth condition, using the Likelihood Ratio Test.

**Table S4.** List of DEGs determined for cork oak phellem samples between WD and WW conditions.

**Table S5.** List of DEGs determined for cork oak inner bark samples between WD and WW conditions.

**Table S6.** List of DEGs determined for cork oak xylem samples between WD and WW conditions.

**Table S7.** Functional GO enrichment analysis of DEGs found for phellem between WD and WW conditions.

**Table S8.** Functional GO enrichment analysis of DEGs found for inner bark between WD and WW conditions.

**Table S9.** Functional GO enrichment analysis of DEGs found for xylem between WD and WW conditions.

## Acknowledgements

We thank Hugo Rodrigues (ITQB NOVA, PT) and Maria Faustino (ITQB NOVA, PT) for participating in sample collection; Mariana Ferreira (ITQB NOVA, PT) for technical assistance in microscopy analysis; and Mafalda Costa and Hugo Matias (ITQB NOVA, PT) for greenhouse support.

## Author contributions

PMB, MMO: conceptualization, funding acquisition, management and supervision; HS: drought assay and Mini-Pam measurements; HS and DL: sample collection, processing and histochemical analysis. DL, RNA extraction and RT-qPCR; PMB and DL: RNA-seq data analysis; PMB, HS and DL: writing the manuscript; all authors edited and reviewed the manuscript.

## Conflict of interest

The authors declare no conflict of interest.

## Funding

This work was supported by FCT - Fundação para a Ciência e a Tecnologia, I.P., through GREEN-IT Bioresources for Sustainability R&D Unit base (DOI: 10.54499/UIDB/04551/2020) and programmatic (DOI: 10.54499/UIDP/04551/2020) funding, LS4FUTURE Associated Laboratory (DOI: 10.54499/LA/P/0087/2020); through Post-Doc contract DL 57/2016/CP1369/CT0029 (awarded to P.M.B.); and the projects CorkREMODEL (DOI: 10.54499/EXPL/ASP-SIL/1190/2021) and SuberInStress (PTDC/BIA-FBT/29704/2017), funded by FCT/MCTES and co-financed by FEDER in the scope of POR Lisboa 2020. This work also was partially supported by PPBI-Portuguese Platform of BioImaging (PPBI-POCI-01-0145-FEDER-022122) co-funded by national funds from OE - ‘Orçamento de Estado’ and by European funds from FEDER - ‘Fundo Europeu de Desenvolvimento Regional’.

## Data availability

Raw RNA-seq reads and read count matrix are available in ArrayExpress (accession E-MTAB-13376) and ENA (ERP151351). The R script used for differential gene expression analysis is available on GitHub (https://github.com/pedro-mb/CorkOakRNAseq). Additional data is available from the corresponding author, Pedro M. Barros, upon request.

## Notes

### Competing Interest Statement

The authors have declared no competing interest.

